# Massively scalable genetic analysis of antibody repertoires

**DOI:** 10.1101/447813

**Authors:** Bryan Briney, Dennis R. Burton

## Abstract

With technical breakthroughs in the throughput and read-length of next-generation sequencing platforms, antibody repertoire sequencing is becoming an increasingly important tool for detailed characterization of the immune response. There is a need for open, scalable software for the genetic analysis of repertoire-scale antibody sequence data. To address this gap, we have developed the ab[x] package of software tools. There are three core components of the ab[x] toolkit, all of which are freely available: abcloud (github.com/briney/abcloud) for deployment and management of computational resources on Amazon’s Elastic Compute Cloud; abstar (github.com/briney/abstar) for pre-processing, germline gene assignment and primary annotation of antibody sequence data; and abutils (github.com/briney/abutils), which provides utilities for interactive downstream analysis of antibody repertoire data.

## Introduction

The human antibody repertoire is vast and complex, with total estimated pre-immune repertoire diversity as high as 10^12^ unique antibody variable regions [1]. This exceptional diversity is accomplished in the heavy chain initially through a process of somatic recombination in which V, D and J gene segments are recombined to form a complete antibody variable region [2]. The fully recombined heavy chain, which also contains a constant region, is then paired with an independently recombined light chain to form a complete antibody. The initial combinatorial diversity is increased by the incorporation of somatic mutations upon antigen stimulation [3,4]. Although early antibody repertoire studies were constrained by low-throughput sequencing technologies, recent advances have resulted in a massive expansion in the scale at which the genetic composition of expressed antibody repertoires can be analysed. Starting in 2009, read-length increases on the 454 GS-FLX sequencer first enabled high-throughput sequencing of full-length antibody variable regions [5–10]. Advanced desktop-scale next-generation sequencing (NGS) platforms, including the Illumina MiSeq and Life Technologies Personal Genome Machine (PGM), can now produce 25-fold higher throughput than the original 454 GS-FLX [11–13]. Despite vastly higher throughput, these desktop sequencers are much more affordable and operate at a much lower cost-per-base, enabling individual laboratories to integrate antibody repertoire sequencing directly into their experimental workflow. Recent read-length increases on Illumina’s HiSeq and anticipated increases on Illumina’s NextSeq and NovaSeq instruments promise an additional throughput increase of several orders of magnitude beyond current desktop sequencers in the near future. The highest throughput sequencers currently available, Illumina’s NovaSeq instruments, produce billions of sequencing reads per run, throughput that approximates the total number of circulating B cells in an adult human. Researchers are on the verge of being able to perform, for the first time, true repertoire-scale genetic analysis of the circulating antibody repertoire.

Unfortunately, as capacity for antibody sequencing has rapidly grown, bioinformatic analysis tools have not kept pace. Utilities for sequence pre-processing (adapter removal, quality trimming, deduplication, merging paired-end reads, etc) have existed for some time [14–19], and the principles behind these tools have been adapted for antibody-specific uses [20–22]. Tools for post-processing (clonal lineage assignment and visualization, repertoire diversity, etc) of antibody repertoire data have also been reported [22–28]. These tools can be assembled into sophisticated antibody repertoire analysis pipelines, but all rely on a single core dependency: V(D)J germline assignment and primary sequence annotation. Several software packages for performing immunoglobulin germline assignment already exist, including IMGT/HighV-QUEST, IgBLAST, iHMMuneAlign, VDJsolver, SoDA, MiXCR, partis, and JoinSolver [23,29–35], but these existing packages contain significant drawbacks that make them unsuitable for processing antibody sequences at repertoire scale. The International Immunogenetics Information System (IMGT) provides germline assignment tools (IMGT/V-QUEST and IMGT/HighV-QUEST) and maintains a gold-standard database of germline gene segments. Unfortunately, although both IMGT/V-QUEST and IMGT/High/V- QUEST are widely used and considered highly accurate, both are closed source and not available as stand-alone software packages in any form. Since the IMGT/V-QUEST and IMGT/HighV-QUEST webservers are limited to 50 or 500,000 sequences per upload, respectively, neither is suitable for processing even the output of current desktop sequencers. JoinSolver shares IMGT/HighV-QUEST’s closed source and webserver limitations. SoDA is closed source and available only as a compiled Windows binary, making it suboptimal for deployment in predominantly Linux-based HPC environments. VDJsolver and iHMMuneAlign are only able to analyse heavy chains and are incapable of properly annotating sequencer- or somatic hypermutation (SHM)-induced insertions and deletions (indels). Although several annotation tools are open source and available for stand-alone use, we note a recurring tendency of IgBLAST, MiXCR and partis to produce incorrect junction annotations. Additionally, we experienced significant alignment accuracy issues with MiXCR, caused by short alignment gaps in somatically mutated antibody sequences, that result in systematic underestimation of mutation frequency and incorrect annotation of indels. Finally, partis is only compatible with legacy versions of Python, making its incorporation into modern Python workflows cumbersome. In light of these limitations, we have developed the ab[x] suite of tools, which contains software tools for launching, configuring and managing clusters of instances on Amazon’s Elastic Compute Cloud (EC2); massively scalable antibody germline assignment and primary sequence annotation; and methods and utilities for interactive downstream analysis. The entire ab[x] suite is freely available and open source under the permissive MIT software license.

## Methods

Full leukopaks (3 blood volumes) were obtained from ten human subjects (Hemacare). Samples were collected at Hemacare’s Southern California donor center. Sample collection was performed under a protocol approved by the Institutional Research Boards of Scripps Research and Hemacare. Informed consent was obtained from each subject. All subjects were healthy, HIV-negative adults between the ages of 18-30 with no reported acute illness in the 14 days prior to leukapheresis. The subject pool was gender balanced and evenly divided between African-American and Caucasian individuals (ethnicity was self-reported; Extended Data Table 1). Immediately upon receipt of the leukopak, peripheral blood mononuclear cells (PBMCs) were purified by gradient centrifugation and cryopreserved.

For each subject, total RNA was separately isolated from 6 aliquots of approximately 5×10^8^ cryopreserved PBMCs (RNeasy Maxi, Qiagen). For each RNA aliquot, antibody genes were amplified in triplicate (18 total samples per subject), with each of the technical replicates processed independently and starting with a separate aliquot of the RNA sample. To minimize the likelihood of cross-contamination between subjects, RT and PCR reactions for each subject were processed in isolation, such that samples from two different subjects were never in proximity during amplification reaction preparation. All amplification primers have been previously described [27]. cDNA synthesis was performed on 11ul of RNA using 10pmol of each primer in a 20ul total reaction (SuperScript III, Thermo Fisher Scientific) using the manufacturer’s protocol and the following thermal cycling program: 55C for 60 minutes, 70C for 15 minutes. Residual primers and dNTPs were degraded enzymatically (ExoSAP-IT, Thermo Fisher Scientific) according to the manufacturer’s protocol. The entire enzyme-treated cDNA synthesis product was used in a 100ul second strand synthesis reaction using 10pmol of each primer (HotStarTaq Plus, Qiagen) using the following thermal cycling protocol: 95C for 5 minutes, 55C for 30 seconds, 72C for 10 minutes. Residual primers and dNTPs were again degraded enzymatically (ExoSAP-IT) and dsDNA was purified using 0.8 volumes of SPRI beads (AmpureXP, Beckman Coulter Genomics) and eluted in 50ul of water. Antibody genes were amplified using 40ul of eluted dsDNA and 10pmol of each primer in a 100ul total reaction volume (HotStarTaq Plus) using the following thermal cycling program: 95C for 5 minutes; 25 cycles of: 95C for 30 seconds, 58C for 30 seconds, 72C for 2 minutes; 72C for 10 minutes. DNA was purified from the PCR reaction product using 0.8 volumes of SPRI beads (AmpureXP) and eluted in 50ul of water. 10ul of the eluted PCR product was used in a final indexing PCR (HotStarTaq Plus) using 10pmol of each primer in 100ul total reaction volume and using the following thermal cycling program: 95C for 5 minutes; 10 cycles of: 95C for 30 seconds, 58C for 30 seconds, 72C for 2 minutes; 72C for 10 minutes. PCR products were purified with 0.7 volumes of SPRI beads (SPRIselect, Beckman Coulter Genomics) and the entire set of samples from a single subject were eluted in a single 120ul volume of water.

SPRI-purified sequencing libraries were initially quantified using fluorometry (Qubit, Thermo Fisher Scientific) before size determination using a bioanalyzer (Agilent 2100). Libraries were re-quantified using qPCR (KAPA Biosystems) before sequencing on an Illumina HiSeq 2500 using 2×250bp Rapid Run chemistry.

## Results

### The ab[x] package of tools for antibody NGS sequence analysis

The tools that comprise the ab[x] package fall into three major areas: 1) deployment, configuration and management of cloud-based compute resources; 2) sequence data pre-processing, germline gene assignment and annotation; and 3) primitives and utilities for interactive analysis of antibody repertoire data. Abstar, which performs germline gene assignment and primary sequence annotation, is at the core of the ab[x] toolkit. Here, we describe abstar’s function and compare the output to existing assignment and annotation utilities.

### Assignment of germline V, D and J genes

Germline gene assignment is performed on recombined antibody sequences using an iterative process, a brief schematic of which is shown in Figure 1. First, the recombined antibody sequence is queried against a BLAST database of germline variable (V) genes using BLASTn (query parameters: word size=11; match reward=1; mismatch penalty=1; gap open=5; gap extend=2; minimum e-value=1) and the 5 highest scoring germline genes are retained [36,37].

**Figure 1.**
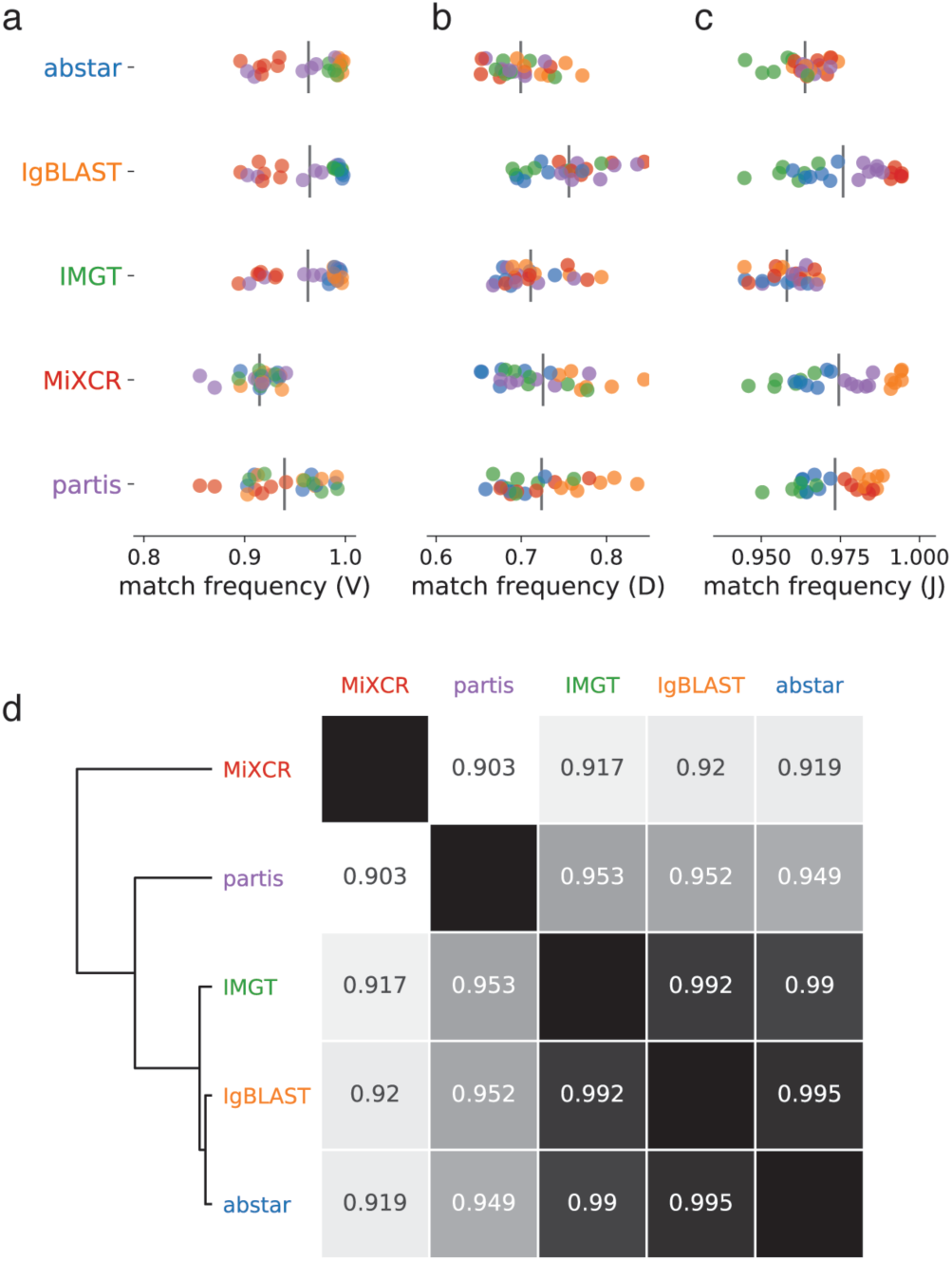
Germline gene segment assignment. Pairwise agreement between assigners on V-gene (A), D-gene (B) or J-gene (C) assignment. Each point represents a single subject, from which 50,000 sequences were analyzed. A clustered distance matrix (D) was computed using the pairwise similarity of each pair of assigners (similarity was the average of the individual similarities for each of the 6 subject sequence sets).

Following variable gene assignment, the portion of the rearranged query sequence that aligns to the germline V gene is removed, the remaining sequence (which represents the junction and joining gene) is queried against a BLAST database of joining (J) genes using BLASTn (query parameters: word size=7; match reward=1; mismatch penalty=-1; gap open=5; gap extend=2) and the 5 highest scoring germline genes are retained. If the rearranged query sequence encodes a heavy chain (determined by checking the top variable gene match), the regions aligning to the germline V and J genes are removed from the rearranged query sequence, the remaining sequence (representing the potential diversity gene and N- and P-addition regions) is aligned to a database of germline diversity (D) genes using Smith-Waterman local alignment (alignment parameters: match reward=3; mismatch penalty=2; gap open penalty=20; gap extend penalty=2) and the 5 highest scoring D genes are retained.

Correlation of germline gene assignment between various antibody analysis programs was tested using a dataset of 300,000 heavy chain sequences isolated from peripheral blood B cells of 6 healthy donors (50,000 sequences per subject). Separately for each subject-derived sequence set, germline gene assignments were performed with abstar, IMGT/HighV-Quest, IgBLAST, partis and MiXCR. With respect to V gene assignment, MiXCR agrees with the other assignment tools least frequently (average of 91% agreement; Figure 1A). Interestingly, the agreement of partis with other assigners was somewhat bimodal, agreeing with IMGT, IgBLAST and abstar about 97% of the time on four subjects but only about 92% of the time on the remaining two subjects (Figure 1A). This effect may be due to partis automatically generating a unique germline database for each subject, meaning the germline database used by partis for any given subject may not precisely match the database used by other assigners. Averaged across all six subjects, IMGT, IgBLAST and abstar behaved the most similarly, with pairwise agreement between these three assigners never less than 99% (Figure 1D). Agreement on J gene assignments was uniformly high, with all programs agreeing 96-98% of the time (Figure 1C). As expected, agreement on the much shorter D gene assignments was far less frequent than was observed for either V or J genes (70-75%; Figure 1B).

### Somatic mutations

BLASTn allows only a limited set of scoring parameters, none of which are optimal for antibody germline gene alignment. Common misalignments include the incorrect splitting of long indel events into multiple shorter indels, likely due to suboptimal gap open/extend penalties, and the improper incorporation of short indels to improve alignment in regions of low homology. To accommodate this issue, abstar realigns the top scoring germline to the rearranged query sequence using a fast Smith-Waterman local alignment algorithm [38] before annotating mutations and SHM-induced indels to ensure the most accurate alignment between the query sequence and the germline variable gene (alignment parameters: match reward=3; mismatch penalty=2; gap open=22; gap extend=1).

We compared the ability of various annotation tools to correctly identify somatic mutations. For each test sequence, the number of nucleotide mutations from germline was determined and compared between assigners. All assigners, with the exception of MiXCR, were in close agreement, typically differing by about 0.5 mutations per sequence (Figure 3A). MiXCR differed from other assigners by an average of approximately 1.5 mutations per sequence. We found this to be due to the incorporation of many short insertions and deletions during germline gene segment alignment which artificially reduce the true mutation frequency. This effect is evident when quantifying the number of frameshift indels (that is, insertions or deletions of a length that is not a multiple of 3). MiXCR alignments produced frameshift indels in over 75% of all sequences, which was dramatically higher than any other assigner (Figure 3B). Partis found frameshift indels in approximately 1.5% of sequences, which was significantly more frequent than either IgBLAST (0.26%) or IMGT (0.36%). Because single nucleotide indels in the variable gene region are virtually always due to sequencing or amplification error, abstar automatically corrects single nucleotide indels, and thus no frameshift indels were annotated by abstar.

**Figure 2.**
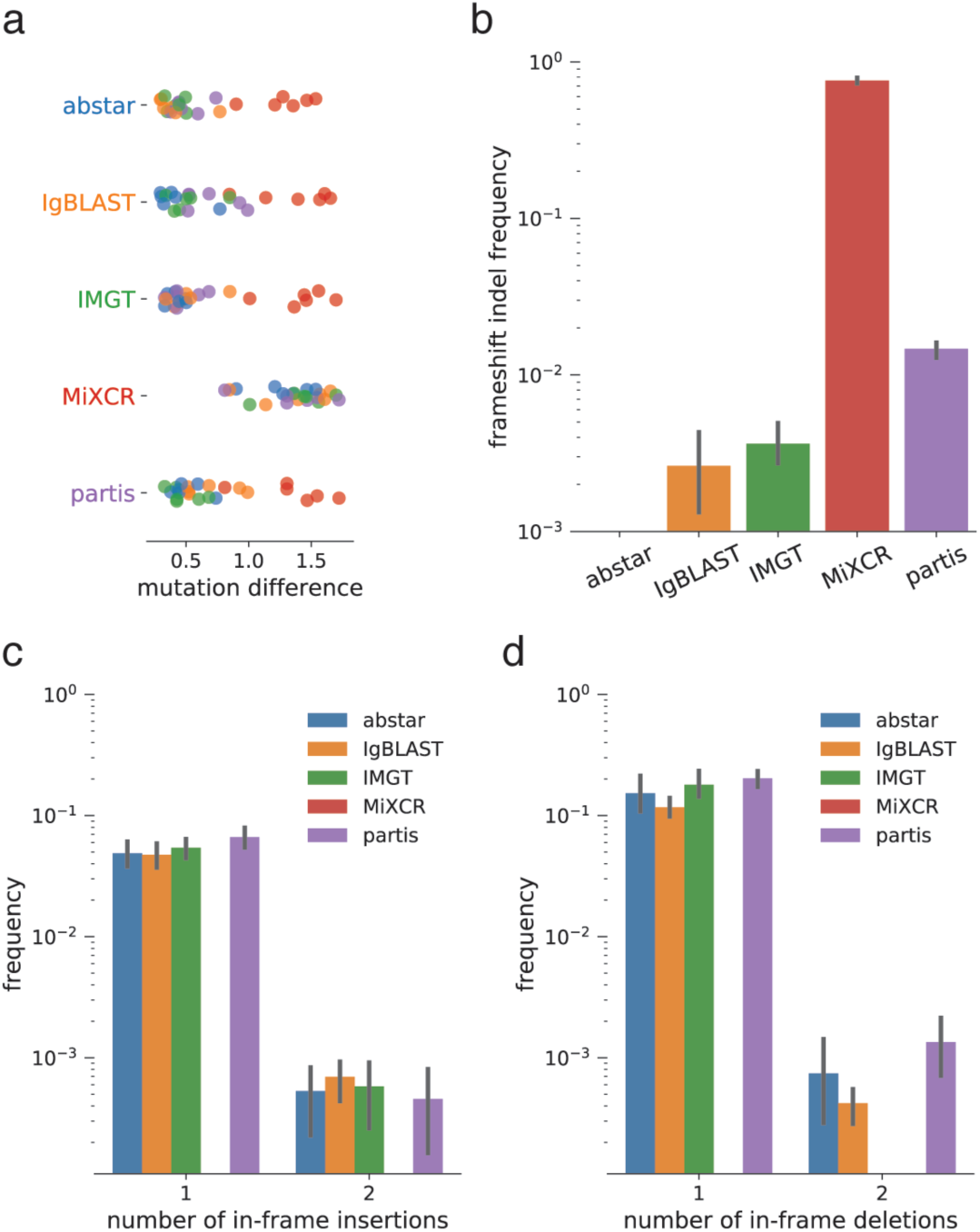
Comparison of nucleotide and indel assignments. (A) For each assigner pair, the absolute value of the difference in mutation count assigned to each sequence was calculated and normalized by the total number of sequences. This value represents the average per-sequence difference in the number of assigned nucleotide mutations. Each point represents the pairwise comparison of the set of sequences derived from a single subject. (B) The number of frameshift insertions or deletions (indels) --that is, insertions or deletions of a length that is not evenly divisible by three -- were quantified for each assigner. Bars represent the average of sequence sets derived from six different subjects, and whiskers represent the 95% confidence interval. The numbers of sequences containing one or two insertions (C) or deletions (D) were calculated. As in (B), bars represent the average frequency of six subjects and whiskers represent the 95% confidence interval. All confidence intervals were calculated in Python using Seaborn (seaborn.pydata.org).

**Figure 3.**
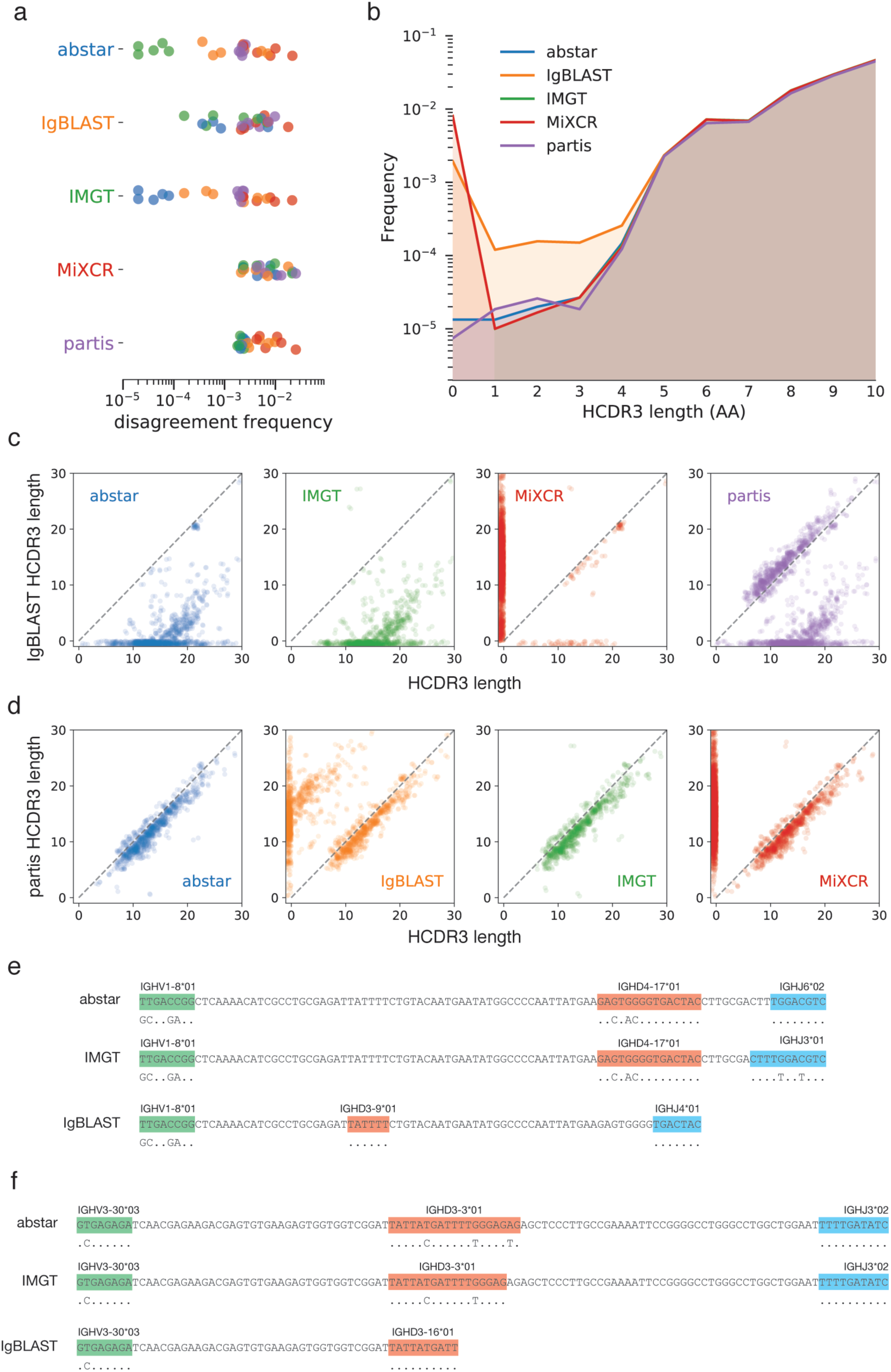
Annotation of antibody junctional regions. (A) The frequency with which pairs of assigners disagree on HCDR3 length were computed. Each point represents the disagreement frequency of a pair of assigners using sequence data derived from a single subject. (B) Frequency which with different assigners assigned short (≤10 amino acids) HCDR3 lengths. (C) Limited to the set of sequences for which IgBLAST disagreed on HCDR3 length with each other assigner, the length assigned by IgBLAST (y-axis) was plotted against the length assigned by the comparison assigner (x-axis). (D) Same as in (C) but using partis as the primary assigner instead of IgBLAST.

Although single nucleotide indels mostly can be attributed to sequencing or amplification error, the SHM process is known to rarely produce relatively short indels [39–41]. These SHM-induced indels are essentially all codon-length, as the low frequency at which they are introduced makes it unlikely that multiple compensatory out-of-frame indels would be accumulated simultaneously. Aside from MiXCR, all assigners were in close agreement about the number of sequences encoding one or two in-frame indels (Figure 3C-D). No sequences in the dataset were assigned more than two in-frame insertions or deletions by any assigner. MiXCR, in contrast, did not assign any in-frame indels, instead splitting codon-length indels into multiple shorter indels, typically only a single nucleotide in length.

### Junction identification

Identification of the antibody junction, which in heavy chains comprises the region between highly conserved cysteine and tryptophan residues, is a core function of antibody annotation utilities. Abstar uses the IMGT numbering system, in which gapped germline gene notation is used to ensure the numbering of conserved positions is consistent in spite of length variation between germline gene segments, to identify the conserved residues flanking the HCDR3 region. IMGT and abstar are in near complete agreement on HCDR3 length assignments (Figure 3A), differing only 21 times in the entire 300,000 sequence test dataset (<0.01%). There appear to be subject-specific differences in the level of agreement between IgBLAST and IMGT or abstar, with three subjects mismatching on approximately 1 in 2000 sequences and the remaining three subjects mismatching roughly 1 in every 200 sequences. This may be due to IgBLAST’s tendency to truncate junctional regions (discussed further below) which produces a larger fraction of very short HCDR3 length assignments (Figure 3B). MiXCR and partis are in agreement with other assigners least often, producing different CDR3 lengths for about 1% of sequences. When partis disagrees with IMGT or abstar, the assigned lengths differ by only one or two amino acids. In contrast, MiXCR appears unable to identify HCDR3 regions in a relatively large fraction of sequences, assigning them a length of zero and causing length differences between MiXCR and other assigners to be quite large (Figure 3B).

In our use of IgBLAST, we have noticed a tendency of the IgBLAST assigner to improperly truncate sequences when analysing antibodies with long HCDR3 regions. One example is shown in Figure 3E, using the anti-HIV antibody PGT145 [42], which encodes an HCDR3 of 33 amino acids. IMGT and abstar agree perfectly on the D gene assignment (IGHV4-17*01) and location. In contrast, IgBLAST assigns a different J gene (IGHJ4*01) to a region identified as the D gene by both other algorithms. This then forces the assignment of a different D gene (IGHV3-9*01) and truncation of the antibody sequence prior to the end of the junction. A second example is another anti-HIV antibody, CAP256-VRC26.08 [43], which encodes an HCDR3 of 37 amino acids. When processed by IgBLAST, the CAP256- VRC26.08 sequence is improperly truncated near the midpoint of the junction (Figure 3F). This results in a different D gene assignment than either IMGT or abstar (IGHD3-16*01 for IgBLAST, IGHD3-3*01 for IMGT and abstar) and no J gene assignment by IgBLAST. Although we initially noted this phenomenon only when analyzing antibody sequences encoding long HCDR3s, based on the CDR3 lengths of sequences for which IgBLAST disagrees with other assigners (Figure 3C), this appears to be a systemic issue that occurs across the CDR3 length distribution.

These alignments (Figure 3E-F) also provide interesting insight into some of the underlying assumptions that result in disagreement between assigners. In the case of PGT145 (Figure 3E), abstar and IMGT differ on the J gene assignment. IMGT prefers the longer alignment (IGHJ3*01) even though the alignment contains two mismatches, however, abstar prefers the identically aligned gene segment (IGHJ6*02) despite shorter overall alignment length. For CAP256-VRC25.08 (Figure 3F), abstar assigns a slightly longer region to the D gene segment, even though extending the D gene alignment region by two positions results in the incorporation of an additional mismatch. IMGT, in contrast, assigns the same D gene but shortens the region to which the D gene is aligned to avoid incorporating an additional mismatch. In both of these cases, it is very difficult to determine which assigner is correct without additional information (perhaps the subject from which PGT145 was isolated expresses a novel IGHJ3 allele which would eliminate one or more of the mismatches in the IMGT alignment, for example). Therefore, it is important to note that agreement between assigners is not necessarily an indication of gene segment assignment correctness; it may simply indicate that the agreeing assigners make similar assumptions when encountering ambiguous cases.

### Distributed processing of large sequence sets

Abstar is natively compatible with Unix-based operating systems (tested on Linux and MacOS), with Windows compatibility provided via Docker. Written in Python, individual ab[x] components are installable via pip, the Python package manager. The ab[x] suite is compatible with Python 2.7 and ≥3.5. While abstar can be run locally or on a single remote compute instance, it is often desirable to distribute the computational load across multiple remote compute instances to allow for annotation of very large sequence datasets that could not be completed on a single computer within a reasonable amount of time. To accomplish this, abstar is tightly integrated with abcloud, which manages single instances or clusters of compute instances on Amazon’s Elastic Compute Cloud (EC2). A schematic representation of an abcloud-configured cluster is shown in Figure 4A.

**Figure 4.**
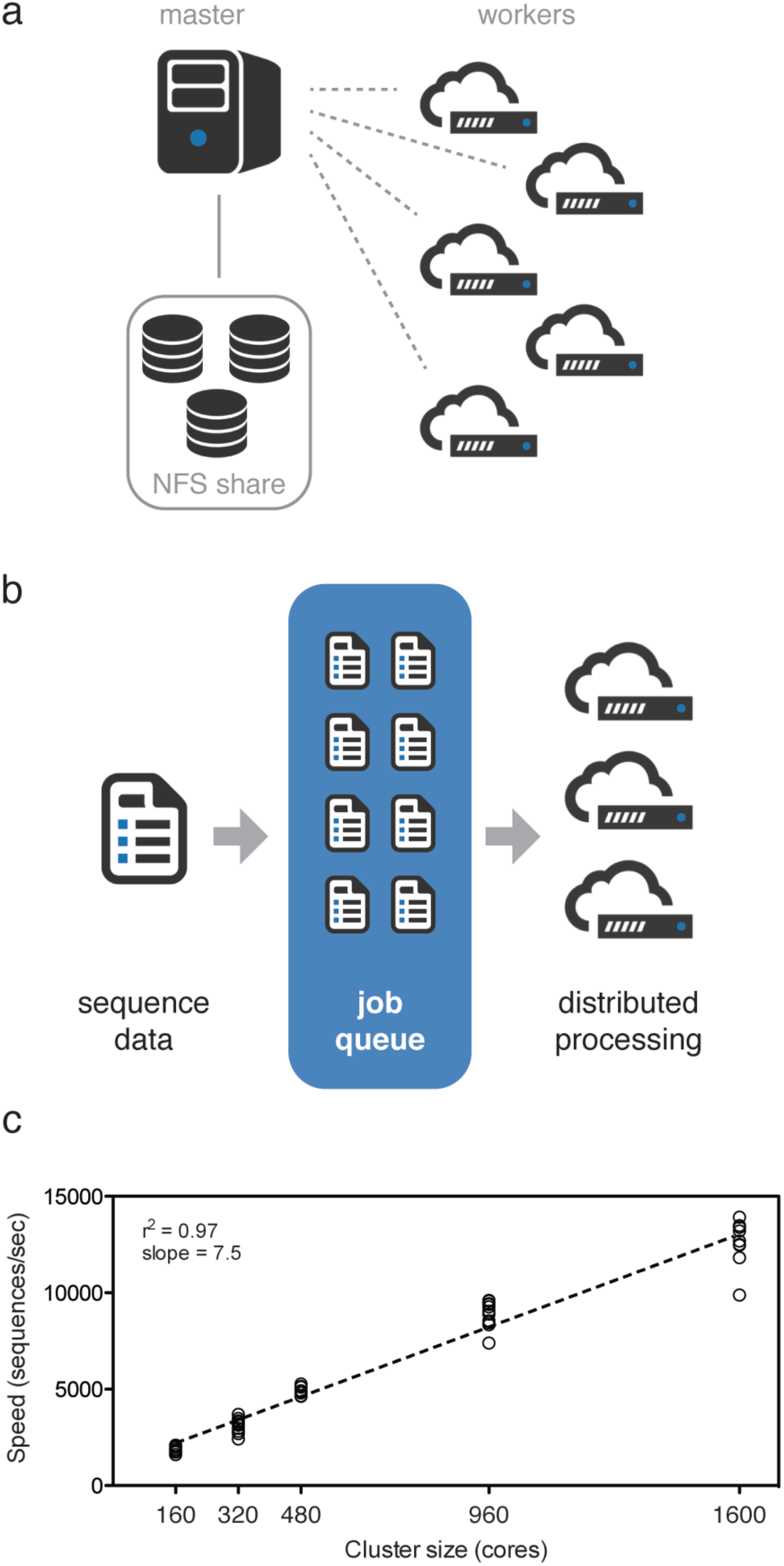
Distributed sequence processing using abcloud and abstar. (A) Schematic of an abstar- managed cluster. The master node shares a large data storage volume with worker nodes using NFS. (B) Schematic of distributed batch processing by abstar. Large input data files are split into smaller “job” files containing approximately 500 sequences. These jobs are submitted to a queue, which distributes and monitors job processing by worker nodes. (C) A test dataset of approximately 25 million human heavy chain sequences (in 11 batches) was processed on five different EC2 clusters. Each cluster consisted of c3.8xlarge worker nodes, with increasing number of instances to produce clusters ranging from 160 to 1600 cores. The processing speed (sequences/second) for each cluster is shown, with individual points representing the different sequence batches.

Abcloud is capable of launching single instances or clusters of instances on EC2 using either reserved or spot pricing. When launching individual instances or clusters of instances, the master node is automatically configured to function as a Jupyter notebook server, for use in interactive downstream analysis. A user-selected number of Elastic Block Storage (EBS) volumes can be attached to the instance and assembled into a redundant array (RAID levels 0, 1, 5, and 10 are supported). When launching a cluster of instances (a single master and one or more workers), the master-attached EBS volume is shared with all worker nodes by default using NFS. Once complete, instances and clusters deployed using abcloud contain all ab[x] tools and required dependencies.

To enable distributed sequence processing on multiple compute nodes, abcloud establishes a Celery distributed task queue (celeryproject.org) with Redis as the message broker and launches Celery worker processes on each of the worker nodes. Briefly, Celery serves as a producer/consumer job queue: the master node (producer) divides the input sequence files into ‘jobs’ containing approximately 500 sequences and deposits them into the queue, while worker nodes (consumers) asynchronously pull jobs from the queue, perform the jobs, and deposit the results into a results queue. A simplified schematic of the queuing process is shown in Figure 4B. The running Celery cluster can be monitored using Flower (github.com/mher/flower). Configured by default on abcloud-managed clusters, Flower provides a convenient web interface for monitoring various aspects of a Celery queue, including completed jobs, remaining queue size, and worker status.

Abstar’s performance was tested using abcloud-deployed EC2 clusters of varying size. The test data was a complete Illumina MiSeq run (approximately 25 million reads) containing 11 pools of antibody heavy chain sequences derived from a single donor. Several clusters were created on EC2 using c3.8xlarge worker instances, with sizes ranging from 160 cores (five c3.8xlarge worker instances) to 1600 cores (fifty c3.8xlarge instances). Performance scales near-linearly with increasing cluster size (Figure 4C). Also of note, because input sequence files are broken down into similarly sized Celery jobs without regard to the size of the input data, memory use increases only marginally with large increases in the amount of input data. Based on these performance numbers and the current spot price of a c3.8xlarge EC2 instance (just under $0.50/hour), annotating a full MiSeq run on a 160-core cluster would be expected to take about 2 hours and cost less than $10.

### Sequence retrieval, preprocessing utilities, and the abstar API

As previously mentioned, several utilities already exist for preprocessing of antibody repertoire sequencing data. However, to ease the creation of software pipelines using abstar as the germline assignment and annotation component, we have incorporated a small set of sequence retrieval and preprocessing utilities into abstar.

Illumina’s family of sequencing instruments are extremely widely used, and reportedly account for more than 90% of all sequencing data ever produced. BaseSpace is Illumina’s cloud-based data storage platform, into which sequencing data may be automatically uploaded in real-time during sequencing runs performed on Illumina’s instruments. To ease the retrieval of sequencing data from BaseSpace, abstar incorporates methods for directly retrieving raw sequencing data from BaseSpace. Using either the BaseSpacePy API (only compatible with Python 2.7) or Illumina’s newer BaseMount utility, abstar users can select a BaseSpace project at runtime rather than an input file or directory of input files. Because BaseSpace is hosted on Amazon Web Services (AWS), data transfer between BaseSpace and EC2 instances is much faster than downloading to a local computer through the BaseSpace web application.

In order to ease the creation of customized sequence analysis pipelines, abstar provides a public API through which core abstar functions can be accessed, as well as methods for standard sequence preprocessing operations. These methods are simply wrappers around commonly used sequence processing tools, with the wrapper providing an easy-to-use interface for pipeline construction using the abstar API. Paired-end sequencing reads can be merged with PANDAseq [17], quality trimming can be performed with sickle (github.com/najoshi/sickle) and adapter trimming can be performed with cutadapt [15]. Run quality statistics can be computed using FASTQC (www.bioinformatics.bbsrc.ac.uk/projects/fastqc). Additional details about the abstar API, including example pipeline scripts, can be found in the abstar documentation (abstar. readthedocs.io).

### Models and methods for interactive downstream analysis

A primary function of the abutils package is to provide a set of core primitives for working with antibody sequence data. This includes object representations of antibody sequences at three levels: sequence, pair, and lineage. At the base is the Sequence object. Sequence objects can be instantiated in a variety of ways, from various string representations of an antibody sequence, from FASTA/Q files, from JSON-formatted abstar output files, or from a MongoDB database populated with abstar-annotated sequence data. If provided with annotated sequence data (either via a JSON file or a MongoDB database), all abstar annotations are accessible directly from the Sequence object using the familiar dictionary-style lookup. Pair objects are built from one or more Sequence objects and are designed to represent paired heavy and light antibody sequences. Lineage objects are assembled from one or more Pair objects and are designed to represent a single clonal lineage. In addition to being used throughout the ab[x] toolkit, these base objects provide an intuitive interface through which downstream analysis may proceed.

Abutils also contains utilities for operations commonly performed during downstream analysis using interactive tools like Jupyter Notebook. As with the preprocessing utilities included in abstar, many of these utilities are simply wrappers around existing tools that allow streamlined interactive use and providing common interfaces. For example, abutils includes a utility for sequence clustering using CD-HIT [45]. In addition to providing an API wrapper for CD-HIT, abutils returns a Cluster object, which has methods for identifying cluster centroids, computing consensus sequences, and visualizing cluster alignment. Sequence alignment is a common process during downstream analysis, and abutils provides utilities for local (Smith-Waterman) and global (Needleman-Wunsch) pairwise sequence alignment. Both forms of pairwise alignment return equivalent Alignment objects which contain a standardized property and method structure, allowing users to switch between either form of pairwise alignment while maintaining code compatibility. Multiple sequence alignment is accomplished with API wrappers for MUSCLE and MAFFT [46,47]. Calculation and visualization of sequence phylogenies is a common method for evaluation maturation of antibody lineages [43,48–50]. Abutils provides methods to estimate phylogentic trees using IgPhyML [51] or, for very large datasets, using FastTree [52]. Finally, abutils provides tools for visualizing phylogenetic trees using the ETE toolkit [53].

## DISCUSSION

At the core of the ab[x] project is the principle that repertoire-scale antibody sequence analysis should be accessible to as many researchers as possible. As capacity for antibody repertoire sequencing rapidly multiplies, data analysis becomes an increasingly difficult technical and financial hurdle. As such we have developed a powerful suite of tools that allow researchers to perform repertoire-scale antibody sequence analysis out-of-the-box. These tools automate and simplify each step of the analysis process, from launching and configuring cloud computing resources to interactive exploratory analysis. Central to the ab[x] toolkit is abstar, which is a best-in-class utility for germline gene assignment and primary sequence annotation. Not only does the ab[x] toolkit provide a solid framework for antibody sequence analysis, but the open source code and permissive licensing (MIT) provide a means for other researchers to adapt and extend ab[x] tools in the future.

## Acknowledgements

The authors would like to thank all study subjects for their participation. This work was supported by the National Institute of Allergy and Infectious Diseases: Center for HIV/AIDS Vaccine Immunology and Immunogen Discovery, UM1AI100663 (DRB); Center for Viral Systems Biology, U19AI135995 (BB), the National Institutes of Health: TSRI Institutional Training Grant, T32-AI007244-32 (BB), the International AIDS Vaccine Initiative (IAVI): Neutralizing Antibody Consortium, SFP1849 (DRB), and the Ragon Institute of MGH, MIT, and Harvard (DRB).

## REFERENCES

1. Alberts B, Johnson A, Lewis J, Raff M, Roberts K, Walter P. The Generation of Antibody Diversity. Garland Science; 2002.

2. Tonegawa S. Somatic generation of antibody diversity. Nature. 1983;302: 575–581.

3. Weigert MG, Cesari IM, Yonkovich SJ, Cohn M. Variability in the lambda light chain sequences of mouse antibody. Nature. 1970;228: 1045–1047.

4. Neuberger MS, Milstein C. Somatic hypermutation. Curr Opin Immunol. 1995;7: 248–254.

5. Boyd SD, Marshall EL, Merker JD, Maniar JM, Zhang LN, Sahaf B, et al. Measurement and clinical monitoring of human lymphocyte clonality by massively parallel VDJ pyrosequencing. Sci Transl Med. 2009;1: 12ra23.

6. Weinstein JA, Jiang N, White RA 3rd, Fisher DS, Quake SR. High-throughput sequencing of the zebrafish antibody repertoire. Science. 2009;324: 807–810.

7. Briney BS, Willis JR, Crowe JE Jr. Human peripheral blood antibodies with long HCDR3s are established primarily at original recombination using a limited subset of germline genes. PLoS One. 2012;7: e36750.

8. Briney BS, Willis JR, McKinney BA, Crowe JE Jr. High-throughput antibody sequencing reveals genetic evidence of global regulation of the naive and memory repertoires that extends across individuals. Genes Immun. Nature Publishing Group; 2012;13: 469.

9. Briney BS, Jr JEC. Secondary mechanisms of diversification in the human antibody repertoire. Front Immunol. 2013;4: 42.

10. Briney BS, Willis JR, Finn JA, McKinney BA, Crowe JE Jr. Tissue-specific expressed antibody variable gene repertoires. PLoS One. 2014;9: e100839.

11. Finn JA, Crowe JE Jr. Impact of new sequencing technologies on studies of the human B cell repertoire. Curr Opin Immunol. 2013;25: 613–618.

12. Georgiou G, Ippolito GC, Beausang J, Busse CE, Wardemann H, Quake SR. The promise and challenge of high-throughput sequencing of the antibody repertoire. Nat Biotechnol. 2014;32: 158–168.

13. Boyd SD, Crowe JE Jr. Deep sequencing and human antibody repertoire analysis. Curr Opin Immunol. 2016;40: 103–109.

14. Edgar RC. Search and clustering orders of magnitude faster than BLAST. Bioinformatics. 2010;26: 2460–2461.

15. Martin M. Cutadapt removes adapter sequences from high-throughput sequencing reads. EMBnet.journal. 2011;17: 10–12.

16. Patel RK, Jain M. NGS QC Toolkit: A Toolkit for Quality Control of Next Generation Sequencing Data. PLoS One. Public Library of Science; 2012;7: e30619.

17. Masella AP, Bartram AK, Truszkowski JM, Brown DG, Neufeld JD. PANDAseq: paired- end assembler for illumina sequences. BMC Bioinformatics. 2012;13: 31.

18. Zhang J, Kobert K, Flouri T, Stamatakis A. PEAR: a fast and accurate Illumina Paired-End reAd mergeR. Bioinformatics. 2014;30: 614–620.

19. Bolger AM, Lohse M, Usadel B. Trimmomatic: a flexible trimmer for Illumina sequence data. Bioinformatics. 2014;30: 2114–2120.

20. Shugay M, Britanova OV, Merzlyak EM, Turchaninova MA, Mamedov IZ, Tuganbaev TR, et al. Towards error-free profiling of immune repertoires. Nat Methods. 2014;11: 653–655.

21. Vander Heiden JA, Yaari G, Uduman M, Stern JNH, O’Connor KC, Hafler DA, et al. pRESTO: a toolkit for processing high-throughput sequencing raw reads of lymphocyte receptor repertoires. Bioinformatics. 2014;30: 1930–1932.

22. Gupta NT, Vander Heiden JA, Uduman M, Gadala-Maria D, Yaari G, Kleinstein SH. Change-O: a toolkit for analyzing large-scale B cell immunoglobulin repertoire sequencing data. Bioinformatics. 2015;31: 3356–3358.

23. Alamyar E, Giudicelli V, Li S, Duroux P, Lefranc M-P. IMGT/HighV-QUEST: the IMGT web portal for immunoglobulin (Ig) or antibody and T cell receptor (TR) analysis from NGS high throughput and deep sequencing. Immunome Res. 2012;8: 26.

24. Moorhouse MJ, van Zessen D, IJspeert H, Hiltemann S, Horsman S, van der Spek PJ, et al. ImmunoGlobulin galaxy (IGGalaxy) for simple determination and quantitation of immunoglobulin heavy chain rearrangements from NGS. BMC Immunol. 2014;15: 59.

25. Safonova Y, Bonissone S, Kurpilyansky E, Starostina E, Lapidus A, Stinson J, et al. IgRepertoireConstructor: a novel algorithm for antibody repertoire construction and immunoproteogenomics analysis. Bioinformatics. 2015;31: i53–61.

26. Cortina-Ceballos B, Godoy-Lozano EE, Sámano-Sánchez H, Aguilar-Salgado A, Velasco-Herrera MDC, Vargas-Chávez C, et al. Reconstructing and mining the B cell repertoire with ImmunediveRsity. MAbs. 2015;7: 516–524.

27. Briney B, Le K, Zhu J, Burton DR. Clonify: unseeded antibody lineage assignment from next-generation sequencing data. Sci Rep. 2016;6: 23901.

28. Kwong PD, Chuang G-Y, DeKosky BJ, Gindin T, Georgiev IS, Lemmin T, et al. Antibodyomics: bioinformatics technologies for understanding B-cell immunity to HIV-1. Immunol Rev. 2017;275: 108–128.

29. Souto-Carneiro MM, Longo NS, Russ DE, Sun H-W, Lipsky PE. Characterization of the human Ig heavy chain antigen binding complementarity determining region 3 using a newly developed software algorithm, JOINSOLVER. J Immunol. 2004;172: 6790–6802.

30. Volpe JM, Cowell LG, Kepler TB. SoDA: implementation of a 3D alignment algorithm for inference of antigen receptor recombinations. Bioinformatics. 2005;22: 438–444.

31. Gaëta BA, Malming HR, Jackson KJL, Bain ME, Wilson P, Collins AM. iHMMune-align: hidden Markov model-based alignment and identification of germline genes in rearranged immunoglobulin gene sequences. Bioinformatics. 2007;23: 1580–1587.

32. Ye J, Ma N, Madden TL, Ostell JM. IgBLAST: an immunoglobulin variable domain sequence analysis tool. Nucleic Acids Res. 2013;41: W34–40.

33. Paciello G, Acquaviva A, Pighi C, Ferrarini A, Macii E, Zamo’ A, et al. VDJSeq-Solver: In Silico V(D)J Recombination Detection Tool. PLoS One. Public Library of Science; 2015;10: e0118192.

34. Bolotin DA, Poslavsky S, Mitrophanov I, Shugay M, Mamedov IZ, Putintseva EV, et al. MiXCR: software for comprehensive adaptive immunity profiling. Nat Methods. Nature Research; 2015;12: 380–381.

35. Ralph DK, Matsen FA 4th. Consistency of VDJ Rearrangement and Substitution Parameters Enables Accurate B Cell Receptor Sequence Annotation. PLoS Comput Biol. journals.plos.org; 2016;12: e1004409.

36. Altschul SF, Gish W, Miller W, Myers EW, Lipman DJ. Basic local alignment search tool. J Mol Biol. 1990;215: 403–410.

37. Camacho C, Coulouris G, Avagyan V, Ma N, Papadopoulos J, Bealer K, et al. BLAST+: architecture and applications. BMC Bioinformatics. 2009;10: 421.

38. Zhao M, Lee W-P, Garrison EP, Marth GT. SSW library: an SIMD Smith-Waterman C/C++ library for use in genomic applications. PLoS One. 2013;8: e82138.

39. de Wildt RM, van Venrooij WJ, Winter G, Hoet RM, Tomlinson IM. Somatic insertions and deletions shape the human antibody repertoire. J Mol Biol. 1999;294: 701–710.

40. Lantto J, Ohlin M. Functional consequences of insertions and deletions in the complementarity-determining regions of human antibodies. J Biol Chem. 2002;277: 45108–45114.

41. Briney BS, Willis JR, Crowe JE Jr. Location and length distribution of somatic hypermutation-associated DNA insertions and deletions reveals regions of antibody structural plasticity. Genes Immun. 2012;13: 523–529.

42. Walker LM, Huber M, Doores KJ, Falkowska E, Pejchal R, Julien J-P, et al. Broad neutralization coverage of HIV by multiple highly potent antibodies. Nature. 2011;477: 466–470.

43. Doria-Rose NA, Schramm CA, Gorman J, Moore PL, Bhiman JN, DeKosky BJ, et al. Developmental pathway for potent V1V2-directed HIV-neutralizing antibodies. Nature. 2014;509: 55–62.

44. Scharf L, Scheid JF, Lee JH, West AP Jr, Chen C, Gao H, et al. Antibody 8ANC195 reveals a site of broad vulnerability on the HIV-1 envelope spike. Cell Rep. 2014;7: 785–795.

45. Li W, Godzik A. Cd-hit: a fast program for clustering and comparing large sets of protein or nucleotide sequences. Bioinformatics. 2006;22: 1658–1659.

46. Edgar RC. MUSCLE: multiple sequence alignment with high accuracy and high throughput. Nucleic Acids Res. 2004;32: 1792–1797.

47. Katoh K, Standley DM. MAFFT multiple sequence alignment software version 7: improvements in performance and usability. Mol Biol Evol. 2013;30: 772–780.

48. Liao H-X, Lynch R, Zhou T, Gao F, Alam SM, Boyd SD, et al. Co-evolution of a broadly neutralizing HIV-1 antibody and founder virus. Nature. 2013;496: 469–476.

49. MacLeod DT, Choi NM, Briney B, Garces F, Ver LS, Landais E, et al. Early Antibody Lineage Diversification and Independent Limb Maturation Lead to Broad HIV-1 Neutralization Targeting the Env High-Mannose Patch. Immunity. 2016;44: 1215–1226.

50. Rogers TF, Goodwin EC, Briney B, Sok D, Beutler N, Strubel A, et al. Zika virus activates de novo and cross-reactive memory B cell responses in dengue-experienced donors. Sci Immunol. 2017;2. doi:10.1126/sciimmunol.aan6809

51. Hoehn KB, Lunter G, Pybus OG. A Phylogenetic Codon Substitution Model for Antibody Lineages. Genetics. 2017;206: 417–427.

52. Price MN, Dehal PS, Arkin AP. FastTree 2 – Approximately Maximum-Likelihood Trees for Large Alignments. PLoS One. Public Library of Science; 2010;5: e9490.

53. Huerta-Cepas J, Dopazo J, Gabaldón T. ETE: a python Environment for Tree Exploration. BMC Bioinformatics. 2010;11: 24.

